# Selenoprotein synthesis is not induced by hepatotoxic drugs

**DOI:** 10.1101/2023.05.12.540527

**Authors:** Sumangala P. Shetty, Dongming He, Paul R. Copeland

## Abstract

**Background and Aims:** Many of the proteins that contain the amino acid selenocysteine are required for optimal defense against cellular stress. As such, one might expect selenoprotein synthesis to persist or be induced upon cellular insult. Because selenocysteine is incorporated by a complex post-transcriptional mechanism, monitoring the transcription of selenoprotein genes is not adequate to understand the regulation of selenoprotein synthesis. We aimed to determine whether selenoprotein synthesis is regulated by the induction of hepatotoxic stress.

**Methods:** We used hepatotropic clinically relevant drugs to evaluate the regulation of selenoprotein synthesis in human hepatocarcinoma cells.

**Results:** We found that two drugs, benzbromarone and sorafenib, caused significant inhibition of selenoprotein synthesis. However, the loss of selenoprotein expression was not specific as total protein synthesis was similarly down-regulated only by benzbromarone and sorafenib.

**Conclusions:** These results allow us to conclude that these hepatotoxins do not induce or preserve selenoprotein synthesis as a protective mechanism.

**Highlights:**

- The treatment of liver cells with hepatotoxic and hepatotropic compounds does not result in increased synthesis of selenoproteins.
- Compounds that induced the canonical oxidative stress response that features NRF2 activation eliminated selenoprotein synthesis.
- The downregulation of selenoproteins was accompanied by general inhibition of protein synthesis.

## 1. Introduction

The trace element selenium is an essential component of 25 human proteins as the amino acid selenocysteine (Sec). Many of these selenoproteins are oxido-reductases that remediate cellular damage due to oxidation. Additionally, these proteins are involved in a myriad of other functions including thyroid hormone production, protein folding and lipid metabolism^1^. Some selenoprotein genes are transcriptionally upregulated in response to stress. For example glutathione peroxidase 2 (GPX2) and thioredoxin reductase-1 (TXNRD1) transcription is upregulated by ∼10 and ∼2-fold, respectively, as a result of concanavalin A induced liver injury in mice^2^. This effect is mediated through the antioxidant response pathway via NRF2 stabilization and binding to promoters with the antioxidant response element (ARE)^3–5^. However, since selenoprotein synthesis is also regulated at the translational level due to the complex process of Sec incorporation^6^, we sought to determine how the translation of selenoprotein mRNAs is affected by drug treatment in a hepatoma cell line model (HepG2). In this study, we used metabolic labeling with radioactive selenium to determine whether a selection of drugs with known hepatotoxicity^7^ or the liver tropism regulate selenoprotein synthesis. We also measured cell viability and NRF2 activation with an ARE reporter. Surprisingly, we found no evidence of induced selenoprotein expression by this group of drugs, but two of them, benzbromarone and sorafenib, significantly downregulated both selenoprotein and total protein synthesis.

## 2. Materials and Methods

### 2.1 Cell culture and reagents

All of the experiments in which endogenous and exogenous selenoprotein expression was assessed were performed in HepG2 cells grown in EMEM (Life Technologies) plus 10% fetal bovine serum (except where indicated). HepG2 cells stably expressing the pARE-T1-Luciferase (ARE-LUC) construct^10,11^ were obtained from A.N. Kong, Rutgers School of Pharmacy. Hepatotoxins and controls used in this study were all purchased from Millipore Sigma: Benzbromarone, Clozapine, Allopurinol, Sorafenib, D-Mannitol, and Ritonavir. Each of them was reconstituted to either 1 mM or 10 mM stock in DMSO.

### 2.2 Metabolic radiolabeling

WT or ARE-LUC expressing HepG2 were grown in either 24-well or 96-well plates to approximately 70% confluence when the medium was changed to serum-free medium supplemented with 100 nM ^75^Se (University of Missouri Research Reactor; specific activity of ∼500 Ci/g) or 50 uCi/ml 35S Met/Cys (Perkin Elmer). For drug or control treatments, the stocks were diluted to the indicated concentrations with serum free media containing 100 nM ^75^Se before being added to cells. After 24 hours of labeling, cells were lysed in NP40 cell lysis buffer (10mM Tris-Cl, pH 7.4, 1% NP40, 1X Roche Complete protease inhibitors) at 4C for 1 hour followed by centrifugation at 18,000 g for 10 mins and supernatant collected as sample for analysis. Samples were diluted with 4X SDS sample buffer (100 mM Tris-HCl pH 6.8, 4% (w/v) SDS, 0.2% (w/v) bromophenol blue and 20% (v/v) glycerol) and boiled at 95 °C for 5 minutes and spun at max speed for 1 minute. Proteins were resolved by 4-15% gradient SDS gels (Biorad). The gels were dried and exposed to a phosphorimage screen for 24 hrs before being developed on the Typhoon FLA7000 (Cytiva).

### 2.3 Cell viability assay

Cell viability was assessed using alamarBlue reagent (Thermo Fisher Scientific). HepG2 cells in 96 well plates were treated with compounds indicated in Figure 2 in 90 μl of serum-free media supplemented with ^75^Se for 6 hours when 10 ul of alamarBlue reagent was added and fluorescence readings were taken (excitation/emission-560/590). A plot of relative fluorescence units vs. drug concentration was used to determine if cell viability was altered.

### 4.4 ARE activation assay

Cells expressing the ARE-LUC construct were treated with the compounds indicated in Figure 1 for 24 hours and then lysed in NP40 lysis buffer and centrifuged at 18,000 g for 10 minutes. 50ul of the supernatant was used to analyze changes in luminescence readings compared to untreated control cells using a plate luminometer (Berthold).

**Fig. 1.**
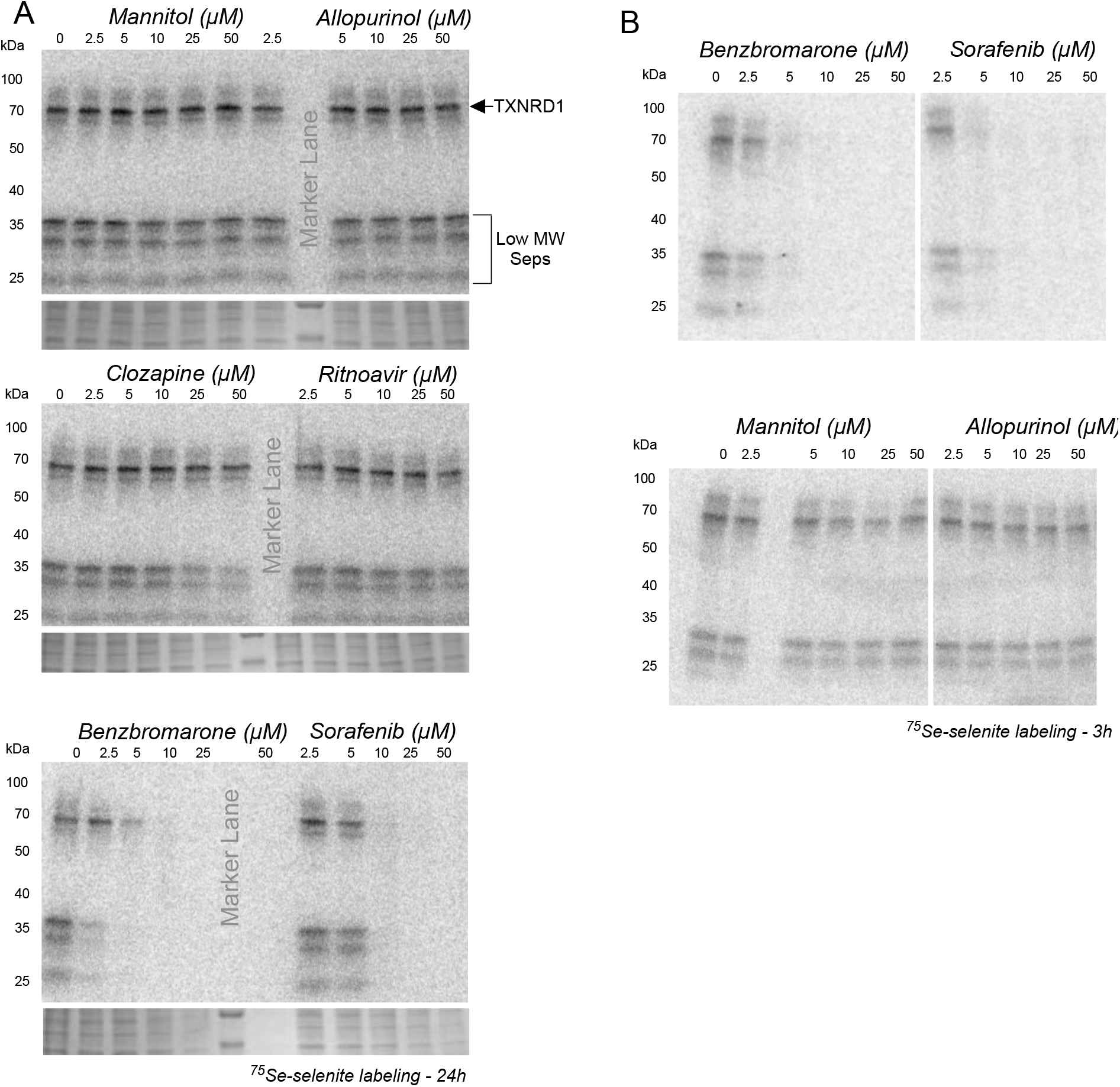
Benzbromarone and sorafenib are potent inhibitors of selenoprotein synthesis in HepG2 cells. A). HepG2 cells were grown for 24 hours in the presence of 100 nM ^75^Se-selenite and the indicated concentrations of drugs. Cells were lysed and cytoplasmic proteins resolved by SDS-PAGE. Gels were stained with Coomassie Blue (lower panels) and radioactive selenoproteins were detected by phosphorimaging (upper panels). B) Same as in A) except metabolic labeling was for 3 hours.

## 3. Results and Discussion

### 3.1 ^75^Se Metabolic labeling reveals downregulation of selenoprotein synthesis

In order to determine the effect of clinically relevant drugs with hepatotoxic potential on selenoprotein synthesis, we metabolically labeled HepG2 cells with ^75^Se-selenite simultaneously with drug treatment ranging from 2.5 to 50 μM for 24 hours. We used three compounds with known hepatotoxicity: 1) the xanthine oxidase inhibitor benzbromarone, 2) the protease inhibitor ritonavir, 3) the antipsychotic clozapine, and one drug with known liver tropism, the chemotherapeutic sorafenib. We also included two controls, D-mannitol and allopurinol, the latter of which is another xanthine oxidase inhibitor that is less toxic^12^. These drugs were chosen for their range of responses in liver, including known activation of the NRF2 response pathway. In addition, liver is a tissue that displays robust selenoprotein production, so regulation at the translational level should be readily apparent.

Selenoprotein synthesis was evaluated by SDS-PAGE of cell lysates followed by Phosphorimaging (Figure 1A). While we saw no evidence of enhanced selenoprotein synthesis, benzbromarone and sorafenib were both strongly inhibitory at 5 and 10 μM, respectively. This inhibition was also apparent after 3 hours of treatment (Figure 1B). It is notable that Benzbromarone seems to differentially inhibit the lower molecular weight (MW) selenoproteins (including the GPXs) relative to the predominant upper molecular weight protein, which is thioredoxin reductase (TXNRD1)^13^. This effect was not observed with Sorafenib treatment, which seems to inhibit selenoprotein synthesis uniformly (Figure 1A, compare 5 μM Benzbromarone to 10 μM Sorafenib). In a similar fashion, the preservation of TXNRD1 synthesis is known to occur under conditions of selenium deficiency^14^, and it is thought to be driven by changes in the amount of methylated Sec tRNA^15^. This finding may suggest that the two drugs inhibit selenoprotein synthesis via two different pathways but further quantitative study is necessary.

### 3.2 Cell death does not drive reduction of selenoprotein synthesis

Although the cells appeared to be healthy in the low inhibitory concentrations of benzbromarone and sorafenib, we sought to determine the effect on viability with the alamarBlue cytotoxicity assay.

HepG2 cells were grown to ∼70% confluence and switched to serum-free medium containing drug concentrations ranging from 2.5 to 25 uM. Viability was assessed at 24 hours post-treatment by alamarBlue staining. Figure 2A shows that none of the cells were sensitive to these drugs at the 3-hour time point but that complete cell death was observed at the 10 μM Benzbromarone concentration after 24 hours of treatment (Figure 2B). We therefore observed no indication that cell death is the primary driver behind the inhibition of selenoprotein synthesis.

**Fig. 2.**
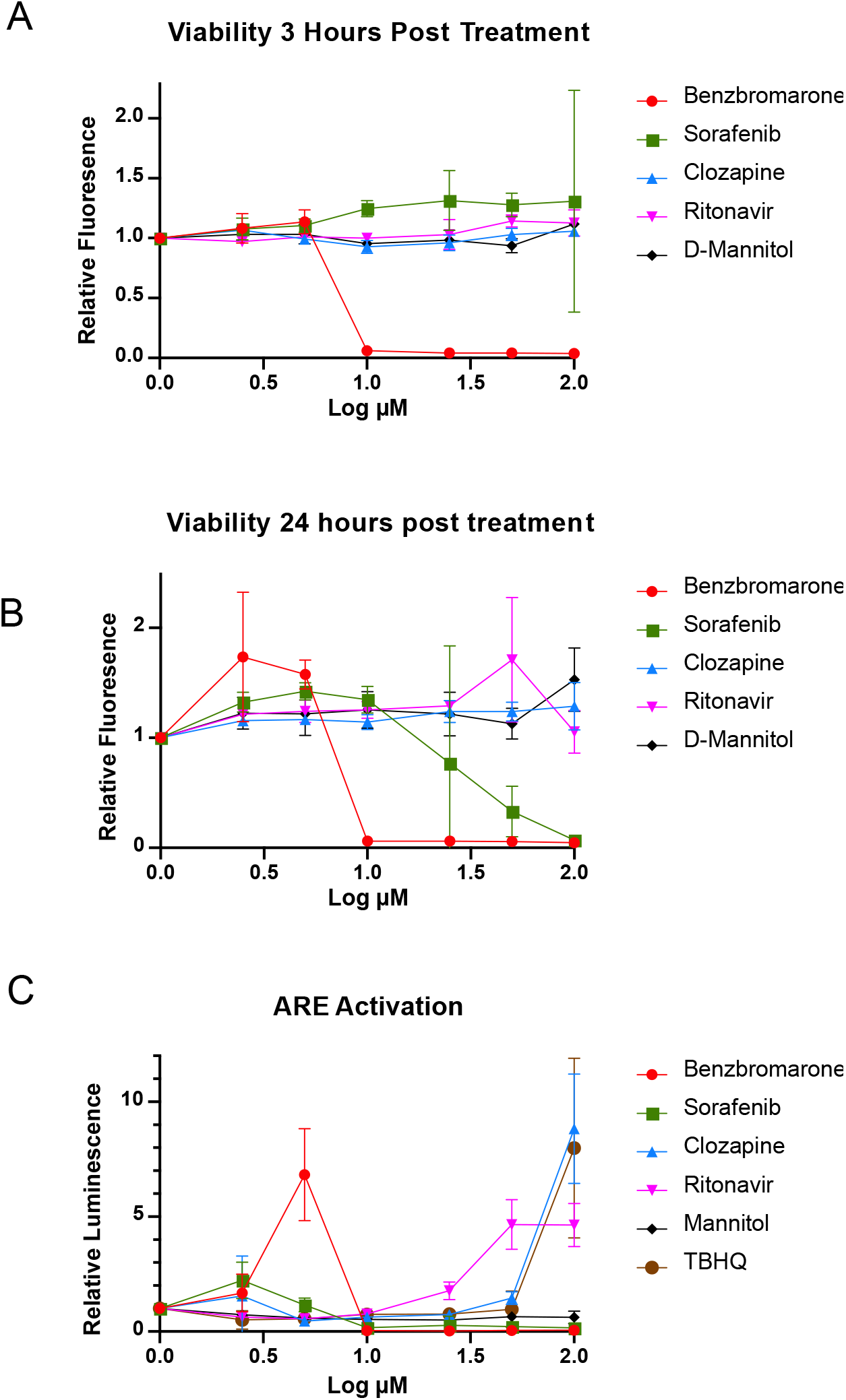
Benzbromarone shows cytotoxicity and ARE activation. A) HepG2 cells were grown in the indicated drug concentrations for 3 hours followed by alamarBlue staining. Fluorescence was plotted versus drug concentration and the results from 3 biological replicates are shown. B) Same as in A except drug treatment was for 24 hours. C) HepG2 cells stably transfected with the ARE-LUC construct were treated with the indicated drug concentrations. TBHQ was included as the postive control for ARE activation. Luminescence was plotted versus drug concentration and the results from 3 biological replicates are shown as the average +/-standard deviation.

### 3.3 NRF2 is activated by benzbromarone

In addition to viability, we also wished to determine the extent to which each of these drugs could activate the antioxidant response element (ARE) pathway driven by NFE2L2 (NRF2). For this experiment we obtained HepG2 cells stably expressing a luciferase construct driven by an ARE-containing promoter^16^. These cells were treated with drug concentrations ranging from 2.5 to 100 μM, which revealed ARE activation at 5 μM for benzbromarone, and 100 μM for the positive control, TBHQ (Figure 2C). No ARE activation was observed for sorafenib or the negative control, D-mannitol. Together these results establish that the loss of selenoprotein synthesis may correlate with NRF2 pathway activation for benzbromarone but not for sorafenib. This is consistent with the selective preservation of TXNRD1 expression during benzbromarone but not sorafenib treatment as shown in Figure 1A.

### 3.4 ^35^S Met labeling reveals a general effect on protein synthesis

To determine whether the inhibition of selenoprotein synthesis is specific, we performed similar experiments using ^35^S Met/Cys labeling to observe total new protein synthesis. For this experiment we trace-labeled the cells with a mixture of ^35^S Met and Cys in the presence of the lower range of benzbromarone, sorafenib and D-mannitol. Figure 3A shows that total protein synthesis appears to be inhibited in a similar fashion as selenoprotein synthesis. Quantitative analysis (Figure 3C) confirms that ^35^S incorporation is inhibited to the same extent as ^75^Se for both benzbromarone and sorafenib.

**Fig. 3.**
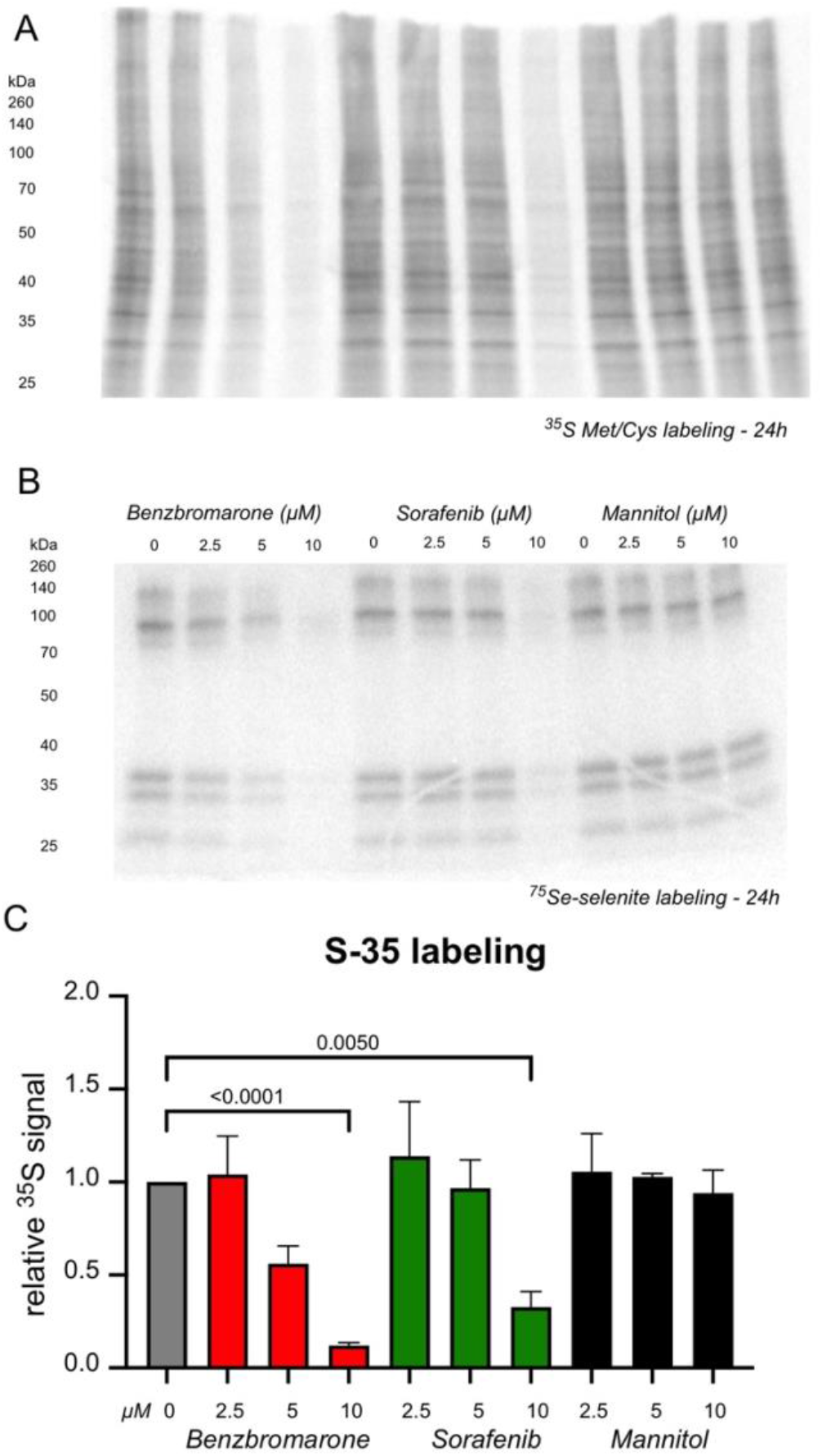
Benzbromarone and sorafenib inhibit total protein synthesis. A) HepG2 cells were grown for 24 hours with the indicated drug concentrations and in the presence of 50 uCi/ml ^35^S Met/Cys mix plus 100 nM cold sodium selenite. Cells were lysed and cytoplasmic proteins resolved by SDS-PAGE. Gels were dried and radioactive proteins were detected by phosphorimaging. B) In parallel with the experiment descrbied in A), cells were labeled with 100 nM ^75^Se-selenite. C) Quantification of total radioactivity in each lane of the 35S labeling gel shown in A). The values were derived from three biological replicates.

Consistent with the prior data, benzbromarone is a more potent inhibitor with substantial loss of signal at 2.5 and 5 μM whereas sorafenib inhibition is only apparent at 10 μM. These results indicate that the inhibition of selenoprotein synthesis is not specific and is likely the result of activation of a stress pathway that inhibits total protein synthesis.

## 4. Conclusions

Here we have demonstrated that the cellular response to the liver tropic drugs tested does not include translational induction of selenoprotein synthesis. Through the use of metabolic labeling with ^75^Se-selenite, we directly evaluated the extent of Sec incorporation after drug treatment, providing a sensitive and specific evaluation of selenoprotein synthesis. By monitoring selenoprotein synthesis with metabolic labeling, we have found that there is no indication of enhanced selenoprotein production under stress conditions induced by hepatotoxic drugs. This finding stands in contrast to the fact that selenoprotein mRNA levels significantly increase upon activation of the NRF2 pathway^2^.This raises the possibility that there may be a disconnect between mRNA and selenoprotein induction as a result of NRF2 activation, which will need further exploration. These results also have implications about the potential side effects of clinically relevant drugs, especially benzbromarone, which is still in use in some parts of the world.

## Abbreviations

(GPX2): glutathione peroxidase 2
(TXNRD1): thioredoxin reductase-1
(ARE): antioxidant response element

## Declarations

No ethical approval was required as this study did not involve human participants or laboratory animals

## Data availability statement

Data associated with this study been deposited into a publicly available repository: BioRxv doi: https://doi.org/10.1101/2023.05.12.540527

## Acknowledgements

Thanks go to Dr. Lauren Aleksunes (Rutgers School of Pharmacy) for initial discussions about hepatotoxins and suggestions. Thanks also to Dr. A.N. Kong (Rutgers School of Pharmacy) for providing the ARE-LUC construct.

